# Compositional and functional differences of the mucosal microbiota along the intestine of healthy individuals

**DOI:** 10.1101/806166

**Authors:** Stefania Vaga, Sunjae Lee, Boyang Ji, Anna Andreasson, Nicholas J Talley, Lars Agréus, Gholamreza Bidkhori, Petia Kovatcheva-Datchary, Junseok Park, Doheon Lee, Gordon Proctor, Dusko Ehrlich, Jens Nielsen, Lars Engstrand, Saeed Shoaie

**Author notes:** Corresponding authors: Jens Nielsen; Lars Engstrand; Saeed Shoaie. These authors share co-first authorship (Stefania Vaga, Sunjae Lee). Abbreviations used in this paper: ARG, antimicrobial resistance gene; CA, caecum; GEM, genome-scale metabolic modelling; MGS, metagenomic species; RE, rectum; TC, transverse colon; TI, terminal ileum.

## Abstract

Gut mucosal microbes evolved closest to the host, developing specialized local communities. There is, however, insufficient knowledge of these communities as most studies have employed sequencing technologies to investigate faecal microbiota. This work used shotgun metagenomics of mucosal biopsies to explore the microbial communities compositions of terminal ileum and large intestine in 5 healthy individuals. Functional annotations and genome-scale metabolic modelling of selected species were then employed to identify local functional enrichments. While faecal metagenomics provided a good approximation of the average gut microbiome composition, mucosal biopsies allowed detecting the subtle variations of local microbial communities. Given their significant enrichment in the mucosal microbiota, we highlight the roles of *Bacteroides* species and describe the antimicrobial resistance biogeography along the intestine. We also detail which species, at which locations, are involved with the tryptophan/indole pathway, whose malfunctioning has been linked to pathologies including inflammatory bowel disease. Our study thus provides invaluable resources for investigating mechanisms connecting gut microbiota and host pathophysiology.

---

Hundreds of thousands of microbial species colonize the mammalian intestine constituting the gut microbiota^1^. This ensemble of species has co-evolved into a complex community in close proximity with the host, developing a symbiosis that provides the host with fundamental functions: protection against pathogens, assimilation of indigestible food, production of essential vitamins, homeostasis maintenance, and immune system development^2, 3^.

The gut microbiota composition is determined by host genetics, diet, lifestyle, ethnicity, and living environment, promoting an important inter-individual variability^4^. The most dominant phyla, in a healthy adult human gut, are Bacteroidetes, Firmicutes, and Actinobacteria^5, 6^. The distribution of specific families, however, is determined by the above-mentioned factors and by local physiological differences, such as pH, oxygen, and nutrients^5^. The gut microbial community composition can therefore rapidly shift in response to both localised and systemic changes. An altered microbiome composition can lead to dysbiosis, which has been shown to affect the onset and progression of several pathologies, such as inflammatory bowel disease, irritable bowel syndrome, diabetes mellitus, obesity, and colorectal cancer^7–9^.

Biogeography of gut microbes is quite heterogeneous^2, 10^. Compared to the gut lumen, the mucus covering the gut mucosa harbours fewer bacteria^5^. The mucus layer lining the epithelium, in particular, is colonised by a unique microbial community, including species such as *Bacteroides fragilis* and *Akkermansia muciniphila*^11, 12^. The importance of A. muciniphila is only starting to emerge in connection with several diseases: it is most beneficial to the host, although an excess has been linked to pathologies such as multiple sclerosis^13^ and Parkinson’s disease^14^. *B. fragilis* has been mostly studied because of its pathogenicity^15, 16^, although it has also been found to provide beneficial effects^17, 18^. Having evolved in closer proximity with the host than any other microbe, they affect the host’s health in several ways that still lack a proper understanding. It is therefore of great interest to further investigate them in their own niche^5, 19^.

The majority of gut microbiome studies have employed faecal sampling for microbiota screening. In order to investigate the gut mucosa microbiota, mucosal biopsies, differently from stool samples, would allow the collection of specific microbial communities. The only studies available on colonic mucosal biopsies have employed 16S ribosomal DNA amplicon sequencing^2, 10, 20–22^ or RNA sequencing^23^, which do not allow a reliable high throughput measurement of the microbial communities^24^.

In this study, we employed shotgun metagenomic sequencing, and a set of functional analysis methods, to investigate the microbiome along the large intestinal mucosa in a healthy subjects’ cohort. The main aim was to investigate if and how this technology can contribute to our understanding of the mucosal microbiota composition and function. We found that, while faecal samples provide a good approximation of the average gut mucosa microbiota, only biopsies could detail subtle but important compositional variations along the intestine. Signature species were identified at each biopsy location, and found, through functional analyses, to affect specific pathways which are known to affect certain physio/pathologic processes.

## Results

To investigate the microbial biodiversity along the large intestine mucosa, we collected faeces samples and, after bowel cleansing, mucosal biopsies, from 5 healthy Swedish adults chosen from a previously published study^25^. Biopsies were taken from 3 locations: terminal ileum (TI), transverse colon (TC), and rectum (RE). As TI was not accessible for subject P1 (Supplementary Table 3), one of her biopsies was taken from caecum (CA) instead (Figure 1A). All samples were prepared for shotgun metagenomics sequencing. We generated, on average, 5.9 million paired-end reads per sample.

**Fig. 1.**
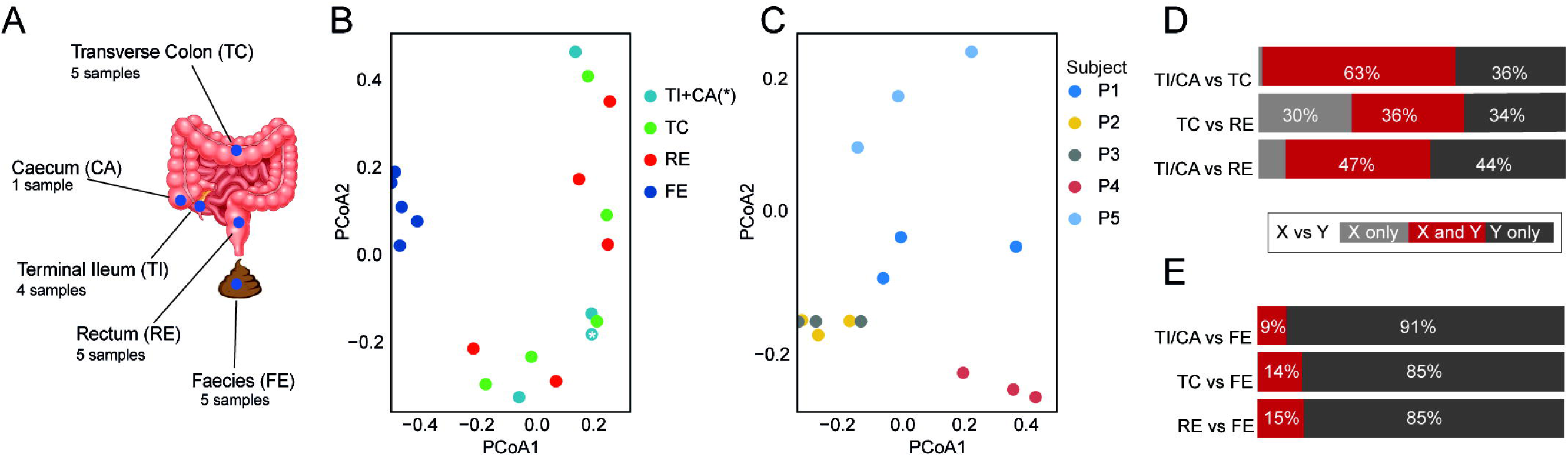
Sampling locations and general overview of the faecal and biopsy-derived metagenomic datasets. **A** For each of the 5 enrolled subjects, gut mucosa biopsies were collected from terminal ileum (TI), transverse colon (TC), and rectum (RE); in one subject, as it was not possible to reach her TI, one biopsy was taken from the adjacent caecum instead (CA). Faeces (FE) were also sampled. **B, C** PCoA plots of the downsized biopsy-faeces dataset **(B)**, colour-coded by sampling-location, and of biopsies data only **(C)**, colour-coded by subject. The only caecal sample of this study is indicated by an asterisk. **D, E** Percentage of shared metagenomic species (in dark red) between biopsy location pairs (D), and between biopsy locations and FE (E).

Whereas faecal samples yielded an average of 5.8 million unique read-counts mapping to the integrated reference catalogue of the human gut microbiome database, only an average of 0.15 million unique read-counts mapped to microbial genes in the biopsy-derived samples. This was due to the majority of reads, in biopsy-derived samples, mapping to human genes (97%). These data were thus used to generate two separately normalised gene-count tables and, correspondingly, MGSs abundance profiles (Supplementary Table 4): one for faeces and one for biopsies. In the next sections, we detail how these datasets were used to investigate the microbial compositions of the biopsies and faeces samples (Supplementary Figure 1).

### Individual microbial uniqueness is stronger than local mucosal microbiota variability

To evaluate the microbial biodiversity in our samples, we first computed their gene- and MGS-richness^26^. We downsized biopsy and faeces datasets together in order to take into account the different sequencing depths across samples (Supplementary Figure 2). In the average, biopsies showed an 82,876 gene-richness, and a 23 MGS-richness. Both these values were about one order of magnitude lower than the corresponding ones for faecal samples (1,093,261 and 259, respectively) (Supplementary Figure 3). Both gene- and MGS-richness showed no significant difference among biopsy locations.

We next looked at how subject- and sampling-location factors contributed to the microbiome diversity. The PCoA plot of the faeces-biopsies dataset showed that the faecal microbiome clustered separately from all the biopsy ones (Figure 1B). While the biopsies microbiomes did not cluster according to their sampling-location (Figure 1B), they did partially cluster based on subjects (Supplementary Figure 4A). This shows that faecal and biopsy microbiomes significantly differ from each other, and that this difference is stronger than individual uniqueness. The PCoA plot of biopsies alone showed a clear clustering based on subjects (Figure 1C), but no clustering based on sampling-location (Supplementary Figure 4B). This suggests that local microbiomes along the length of the large intestine mucosa have subtle differences, which are overcome by individual variability.

To better interpret these findings, we computed the number of shared MGSs between samples (Figure 1, D and E), and between subjects (Supplementary Figure 5). We observed a considerable MGS overlap between biopsy locations (Figure 1D), especially between TI/CA and TC (63%). On the other hand, Figure 1E shows that, whereas almost all the MGSs detected in biopsies were also detected in faeces (only one MGSs detected in TC, *Acetobacter*, was not detected in faeces), the strong separation between faeces and biopsies (Figure 1B) was due to the much higher faecal microbial richness. Within all the MGSs detected in any biopsy-microbiome, 30% were shared by at least two subjects, (Supplementary Figure 5), while this percentage, for the faecal microbiome, reaches 54% (Supplementary Figure 5). This indicates that individual variability is higher in biopsies than in faeces. These results show that, although biodiversity was higher in faeces, since all the MGSs found in biopsies were detected in faeces as well, faecal sampling provides a good approximation of the average gut mucosa microbiota.

### The biopsy-derived gut mucosal microbiomes offer a detailed insight into the biogeography of the large intestine

We next performed a taxonomic analysis of biopsies and faecal microbiomes, in order to assess which MGSs set the two groups apart, and which ones characterize specific biopsy locations. We first computed the total number of MGS-reads belonging to each phylum/class. Figure 2 shows all nine detected phyla (Figure 2A) and the top-ten classes (Figure 2B), sorted from the most to the least abundant in biopsies. Phylum Firmicutes was the dominant one, both in faeces and in biopsies, and it was mostly comprised of Clostridia. The phylum Bacteroidetes was the second most represented in biopsies (all belonging to class Bacteroides), whereas it was third in faeces after Actinobacteria (Supplementary Figure 6). These results are in agreement with current knowledge^5^.

**Fig. 2.**
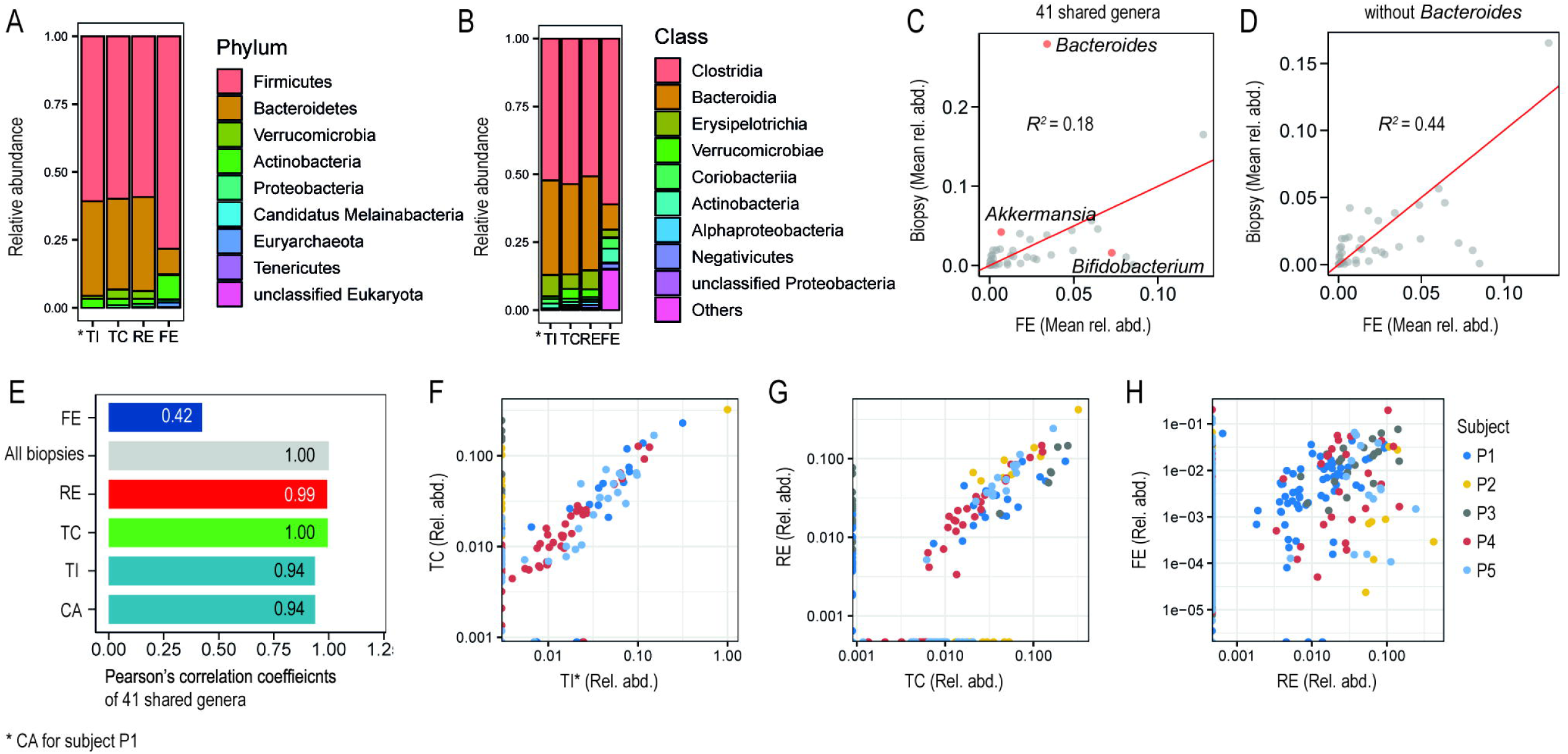
Taxonomy of metagenomics species (MGSs) from terminal ileum/caecum (TI*), transverse colon (TC), rectum (RE), and faeces (FE); n=5 for each group. **A, B** Top 10 most highly abundant phyla **(A)** and classes **(B)** in the large intestine; phyla/classes are sorted, in the legends, from the most to the least abundant in all three biopsy-locations. The corresponding faecal values are also plotted for comparison, in the same order. **C, D** Scatter-plots of mean relative abundances of all 41 genera shared by biopsies and FE **(C)**, and of the same genera except Bacteroides **(D)**. **E** Pearson’s linear correlation coefficients between all biopsies and each sampling location. **F-H** Scatter-plots of the genera relative abundances of three sample-pairs: TI* versus TC **(F)**, TC versus RE **(G)**, and RE versus FE **(H)**; data points are colour-coded by subject.

We next compared the microbiome of all biopsies as a group to the faeces one. To this purpose, we plotted the 41 genera that were shared by these two groups (Figure 2C). The correlation between faeces and biopsies was low (R^2^=0.18) because of the genus *Bacteroides*, which was significantly enriched in biopsies only (Figure 2C). The removal of this genus from the shared genera resulted in a significantly higher correlation between biopsies and faeces (R^2^=0.44; Figure 2D). Pearson’s linear correlation of all shared 41 genera, including *Bacteroides*, between single biopsy locations, was extremely high (R>0.94; Figure 2, E-G), whereas RE and faeces had a much lower correlation (R=0.42; Figure 2H). While these results (Figure 2, E-H) agree with our previous observations, they also show that all the subjects equally contribute to all correlations. We can therefore conclude that the most relevant discrimination factor between faeces and biopsies is genus *Bacteroides*.

Intrigued by the high *Bacteroides* enrichment in all biopsy microbiomes, we looked at which *Bacteroides* species were detected in each biopsy location. Known to be dominant in the gut microbiome^27^, *B. vulgatus* was detected in all samples. Several *Bacteroides* showed specifically high local relative abundances. *B. vulgatus*, *B. thetaiotaomicron*, *B. uniformis*, and *B. caccae* were highest in TI, *B. faecis* in TC, while *B. dorei* and *B. nordii* were only detected in RE. We also observed a decreasing gradient distribution of most of the dominant *Bacteroides* of TI along the large intestine, with the exception of *B. caccae* (Supplementary Figure 7).

The most enriched genus in the faecal microbiota was the carbohydrate-fermenting genus *Bifidobacterium* (phylum Actinobacteria) (Figure 2C). *B. adolescentis* and *B. longum* were detected in the faecal samples of almost all subjects (Supplementary Figure 8). *B. adolescentis*, particularly, displayed an increasing gradient distribution along the large intestine, although its detection in biopsies was significantly lower than in faeces (Supplementary Figure 8).

Our results support our hypothesis that, with the exception of *Bacteroides*, faeces metagenomics provides a good approximation of the average mucosal microbiota. Faecal metagenomics, however, lacks the additional dimension provided by biopsies: a measure of the subtle changes MGSs undergo along the intestine. Since certain pathologies, such as Crohn’s disease, are known to develop at specific locations along the intestine^28–30^, the ability to detect a pathological dysbiosis at such locations is likely to be fundamental for early diagnosis.

### Antimicrobial resistance genes distribution along the large intestine

As the gut microbiota is a reservoir of antimicrobial resistance genes (ARGs)^31^, we next investigated how microbial communities along the intestine contribute to antibiotic resistance. We found that TI was less ARG-rich than both TC and RE, while faeces were significantly richer than any biopsy location (Supplementary Figure 9). This indicates that TI microbiome may be more susceptible to antimicrobials than those from other locations.

We then looked at how the resistomes of this study correlate with one another (Figure 3A, Supplementary Figure 10, Supplementary Figure 11). The faecal resistome had a low correlation both with all biopsies grouped together (R=0.56) (Figure 3A) and with each single biopsy location (Figure 3C). Biopsies’ resistomes, on the other hand, had very high correlations with each other (R>0.85). These results show that biopsies, besides having very similar microbiota (Supplementary Figure 11), had highly overlapping resistomes as well.

**Fig. 3.**
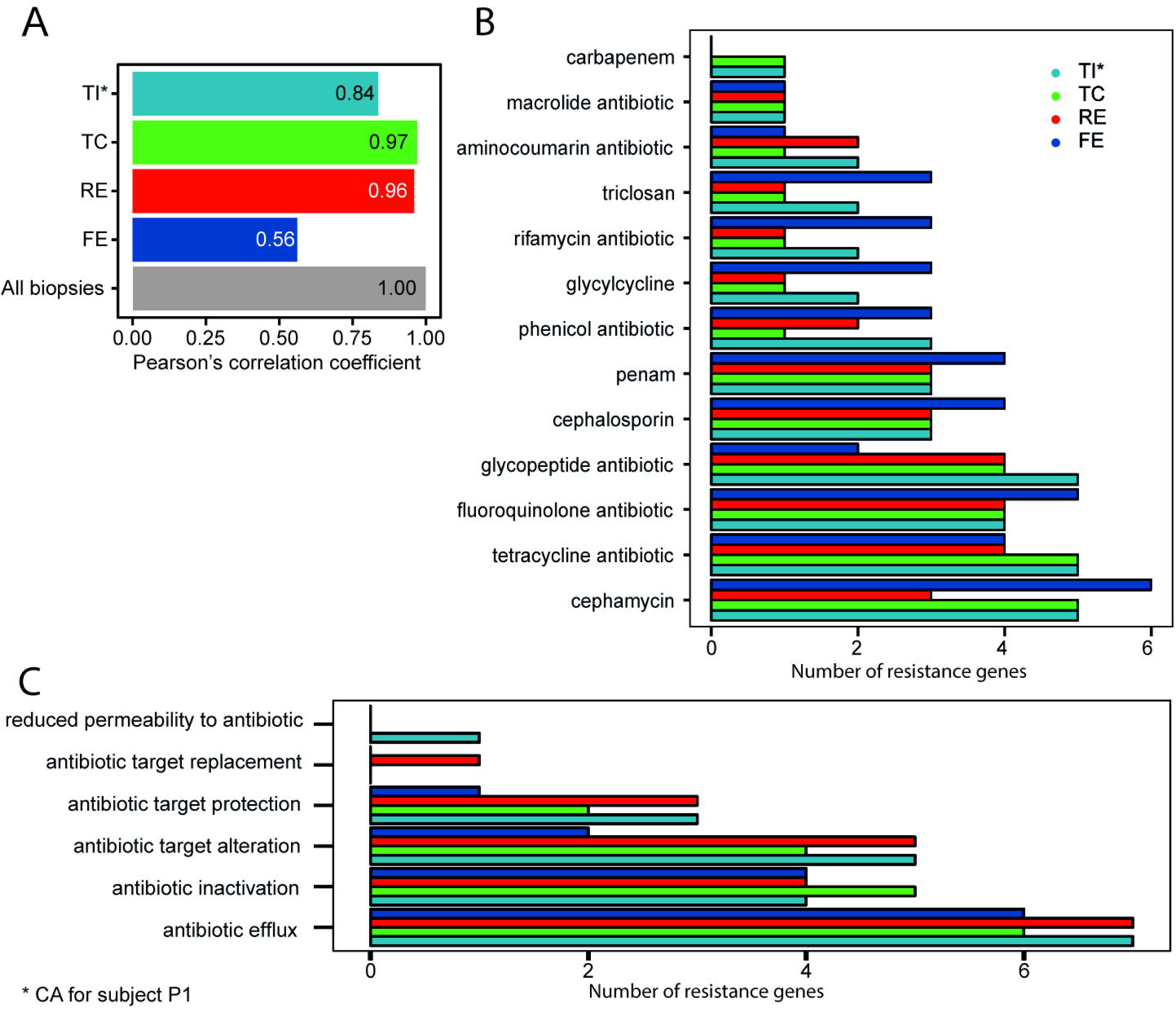
Analysis of antimicrobial resistance genes (ARGs) across biopsy locations (terminal ileum plus caecum (TI*), transverse colon (TC), and rectum (RE)), and in faeces (FE) samples (n=5 for each group). **A** Pearson’s linear correlation coefficients between ARGs relative abundances of all biopsies and each sampling location. **B, C** Differential enrichment of drug classes **(B)**, or of resistance mechanisms **(C)**.

We next investigated the differentially enriched drug classes (Figure 3B) and resistance mechanisms (Figure 3C). The drug classes with the highest number of ARGs were cephamycin, tetracycline, fluoroquinolone, and glycopeptide antibiotics, particularly in TI and TC (Figure 3B). The mechanisms with the highest number of ARGs were *Antibiotic efflux, Antibiotic inactivation*, and *Antibiotic target alteration* (Figure 3C), with particular biopsy locations enrichments. Specifically, *Antibiotic efflux* and *Antibiotic target* alteration were more enriched in TI and RE, while *Antibiotic inactivation* was more enriched in TC. Interestingly, some mechanisms were exclusively represented in one location only, such as *Antibiotic target replacement* in RE, and *Reduced permeability to antibiotic* in TI.

### Functional analysis by KEGG orthology, antiSMASH, and genome-scale metabolic models

To gain a deeper insight into the roles played by locally dominant species along the gut mucosa, we employed a set of functional analysis tools. We first measured how each sample was enriched in specific molecular functional orthologs by assigning KEGG orthologies^32^ (https://www.genome.jp/kegg/ko.html) to genes detected at each biopsy location. Two KEGG orthology pathways were found to be significantly enriched (p-value<0.05) at specific biopsy locations (Supplementary Figure 12): carotenoid biosynthesis and oxidative phosphorylation. The second was enriched in TI, because of locally dominant bacteria such as *Bacteroides vulgatus* and *Bacteroides uniformis*, while carotenoid biosynthesis was enriched in TC due to *Bacteroidetes vulgatus*, *Akkermansia muciniphila*, *Faecalibacterium prausnitzii*, *Parabacteroides* distasonis, among others. As carotenoids have been shown to play a protective role in the human gut by regulating immunoglobulin A (IgA) production^33^, we speculate these microbes may play a role not only in carotenoid synthesis but also in regulating and preserving the gut immune system.

The antiSMASH database^34^ was next employed to annotate the faecal and the biopsy-derived data, in order to find which secondary metabolites (SMs) are preferentially secreted by the microbes found in a specific sample-type. We found a significant enrichment of such metabolites in the faeces samples, compared to all the biopsy locations, due to the higher number of microbial species detected in faeces (Supplementary Figure 13A). Within biopsies, TC was the most enriched location. Nine SMs were enriched in all biopsy locations, and an additional five were only detected in TC and RE (Supplementary Figure 13B). Among these, we found anti-bacterial SMs resorcinol and bacteriocin (secreted by *B. dorei*, *B. faecis*, *B. massiliensis*, and *B. cellulosilyticus*, among others), and aryl polyene (secreted by *Akkermansia muciniphila*, and by most *Bacteroides* species), which provides bacteria with protection from oxidative stress similarly to carotenoids^35^.

We then employed genome-scale metabolic modelling (GEM) to investigate how locally-enriched microbes contribute to the metabolism of the gut microbiota and of the host. Specifically, we constructed and simulated GEMs of all the *Bacteroides* detected in biopsies, and of all the species that were most highly enriched at a single biopsy location, in at least two subjects (Supplementary Figure 14).

**Fig. 4.**
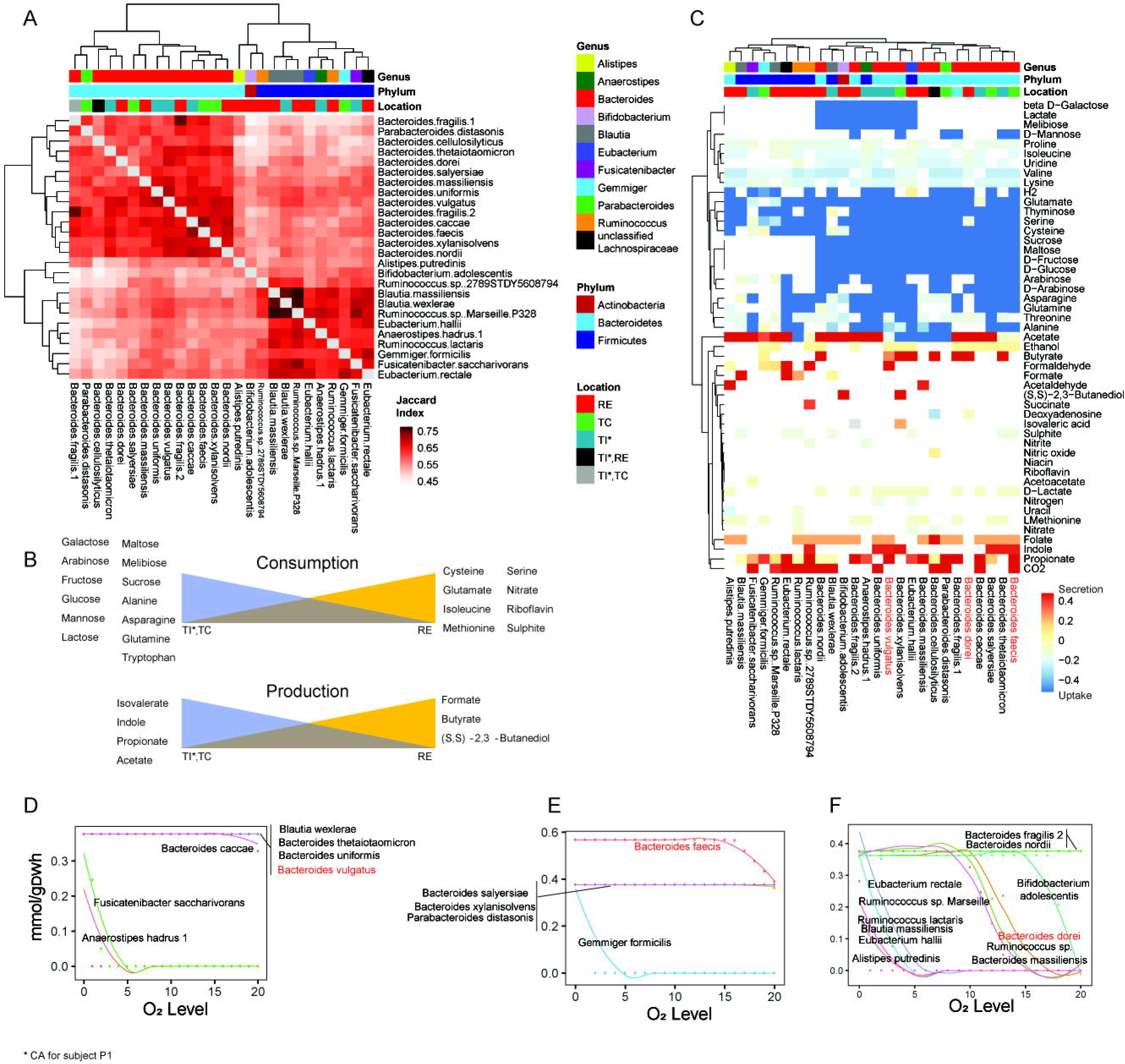
Simulation results of the genome-scale metabolic models of all *Bacteroides* and *Bifidobacterium* species detected in biopsy samples, plus the most highly enriched species at one biopsy location only in at least two patients (Supplementary Figure 10; Supplementary Table 1). Results are reported by biopsy location: terminal ileum plus caecum (TI*), transverse colon (TC), and rectum (RE). **A** Jaccard index for each modelled species. **B, C** Biopsy location-based summation of the secretion of the main bacterial metabolites. **D** Uptake and secretion of the metabolites showing a variation across the modelled species. **E-G** Growth rate plots of the modelled species as a function of environmental oxygen level for TI **(E)**, TC **(F)**, and RE **(G)**. The most enriched *Bacteroides* of the three biopsy locations are highlighted in red (D-G).

The Jaccard similarity analysis between the simulation results of all modelled species was in agreement with our former observations: there was no marked difference between biopsy locations (Figure 4A). However, several metabolites were especially, although not exclusively, produced/secreted at certain biopsy locations because of locally-enriched species (Figure 4, B-C). In particular, we report a gradual shift from sugar to amino acid consuming microbes (Figure 4B). As expected^36^, acetate secretion (highest in TI) was greater than propionate’s, which was greater than butyrate’s (Supplementary Table 2). Formate and butyrate production was highest in RE (Figure 4B), where *Ruminococcus lactaris, Eubacterium* rectale, and *B. nordii* were enriched (Figure 4C), while propionate production was highest in TI (Figure 4B), which had higher concentrations of *B. caccae*, *B. vulgatus*, and *B. fragilis* (Figure 4C).

Our simulations yielded novel interesting insights into several metabolic pathways whose relevance to health or disease has already been assessed, but whose mechanisms are still largely obscure. In particular, we observed significantly enriched indole secretion and L-tryptophan (Trp) consumption in TI and TC (Figure 4C), mainly due to *B. thetaiotaomicron*, *B. uniformis*, *B. vulgatus*, *B. faecis*, and *B. xylanisolvens*. Trp is an essential amino acid involved in several important functions, mostly connected with the host-microbiome interaction. As a fraction of Trp is known to be metabolised into indole by the gut microbiota^37^, our results show that this process mainly happens in TI and TC (Figure 4C).

Finally, by simulating the growth of the modelled species with varying oxygen concentrations, we observed different responses from locally dominant *Bacteroides*. In particular, *B. vulgatus* (dominant in TI) was shown to have a high oxygen-resistance (Figure 4D), whereas *B. faecis* (dominant in TC) showed a lower one (Figure 4E), and *B. dorei* (dominant in RE) was quite oxygen-sensitive (Figure 4F).

## Discussion

The current best strategy to study gut mucosal microbial communities is to sample them through biopsies and to employ a shotgun sequencing technology to measure their microbial species composition. This study was designed to explore the advantages and limitations of using shotgun metagenomics and gut mucosal biopsy samples to quantify local microbiomes along the intestine of healthy subjects.

We show that faecal samples are richer in biodiversity. This is due, at least partially, to the expected lower number of reads mapping to microbial genes in biopsy-derived samples, as these contain higher amounts of human DNA. We also found important individual differences, particularly in the biopsy microbiota, less so in the faecal ones. Biodiversity was, therefore, the main discriminating factor between faeces and biopsies microbiota, followed by individual variability, while biopsy sampling-location offered the least discriminating power across biopsies. Although not significantly different in composition, different gut locations vary in species concentrations: the same species may be found in most or all the biopsy locations, but with a different prevalence within each local community. Such variations are of crucial interest, as they may affect the host metabolism and homeostasis, or the onset and development of certain diseases where an altered composition of local communities is linked with such processes.

Despite the richer faecal biodiversity, faecal microbiota included almost all the MGSs found in mucosal biopsies. Additionally, faecal and biopsy microbiomes correlate with each other very well, with the exception of *Bacteroides*. This suggests that faeces metagenomics provides a good approximation of the average gut mucosal microbiota composition, in health.

Both faeces and biopsies were particularly rich in Firmicutes, a phylum known to be dominant in the gut microbiota^5^. Several microbial species belonging to genus *Bacteroides* were, however, significantly enriched in biopsies only, whereas species from genus *Bifidobacterium* were mainly enriched in faeces. *Bacteroides* are known as obligate anaerobic, bile-resistant bacteria, and one of the dominant genera in the gut^38^. Although they often behave as commensal organisms, some of their features allow them to turn into pathogens^38, 39^. They are among the rare prokaryotes provided with membrane sphingolipids, which are believed to enhance their ability to detect and cope with an unstable environment^40^. Furthermore, *Bacteroides*’ outer membrane comprises lipopolysaccharides^41^; although they usually trigger the host immune response, *Bacteroides* species-specific lipopolysaccharides structural differences prevent that^42^. Overall, *Bacteroides* are equipped with a unique interface with the host. Besides providing them with the capability to quickly detect and react to environmental changes, it also allows them to affect and modulate the host immune system^42^. Among the most enriched *Bacteroides*, we showed *B. thetaiotaomicron* and *B. fragilis* to gradually decrease from TI to RE, while *B. faecis* was significantly enriched in TC. Being among the best-studied *Bacteroides*, they are known as flexible foragers, capable of adapting to changes in microbial composition and glycan availability^43, 44^. Their enrichment may, therefore, have been caused by the depletion of other genera caused by bowel cleansing.

We next performed a set of functional analyses to investigate the roles played by locally dominant species. First, we explored the local enrichment of antibiotic resistance genes. The higher ARG-richness found in faeces was due to the greater MGS-richness of these samples. Interestingly, the number of resistance genes corresponding to several resistance mechanisms was higher in some biopsy locations, particularly TI and RE, than in faecal samples. In particular, although TI had the lowest gene/MGS-richness, and no unique MGSs of its own, it had the highest number of significantly changing ARGs. Some ARGs may exist as a result of acquired resistance to antibiotic use. We next performed a KEGG orthology analysis, which resulted in two pathways being significantly enriched. Of these, the first was carotenoid biosynthesis, which was enriched in TC because of the local prevalence of a few microbes, including *Bacteroidetes vulgatus* and *Akkermansia muciniphila*. The carotenoid biosynthesis pathway is responsible for producing beta-carotene, a precursor for retinol metabolism, and vitamin A. It has been shown that carotenoids play a protective role on the human gut by regulating IgA production^33^, and that retinol deficiency specifically affects *Bacteroides vulgatus’* growth in the gut^45^. Our data allows us to speculate that microbes possessing genes from this pathway, and locally enriched in TC, may play an important role in the retinol turnover of the gut, and in regulating IgA production.

GEM modelling was used to investigate the secretion/uptake of fermentation metabolites by the most enriched microbial species at each biopsy location. The simulations of our models provided new insights into partially known metabolic mechanisms involving microbial species which were found, in this work, to be dominant at specific locations only. We observed, for instance, a gradual shift, along the intestine, from sugar to amino acid consuming microbes, and we reported unprecedented oxygen-resistance details of specific *Bacteroides* species. Of particular interest was the discovery that Trp consumption and indole production were both significantly increased in TI and TC. Trp is an essential amino acid that plays fundamental roles in regulating nitrogen balance, gut immune system homeostasis, and serotonin production^46, 47^. A fraction of Trp is metabolised to indole and its derivatives by the gut microbiota, consistently with our simulation results^46^. Several diseases have been linked with reduced efficiency of this mechanism^47, 48^. In particular, inflammatory bowel disease has been associated with decreased serotonin production and Trp imbalance^49^. *Bacteroides* are among the few known microbes possessing Trp metabolism catalytic enzymes^47^. The main *Bacteroides* found in TI and TC that, according to our GEM simulation, were responsible for Trp consumption and indole secretion, were *B. thetaiotaomicron*, *B. uniformis*, *B. vulgatus*, *B. faecis*, and *B. xylanisolvens*. This novel information is extremely valuable, particularly as the impact of this mechanism is only starting to be uncovered in connection with a growing number of diseases. The *Bacteroides* species we report here in connection with Trp/indole metabolism will thus require further targeted investigation, as that could help prevent and treat such pathologies. As the results derived from our GEM simulation were obtained *in silico*, further targeted experiments will be required in order to validate them.

One of the main limitations of this study was the small number of subjects that could be enrolled. Very strict inclusion criteria were applied to ensure that all the included subjects were healthy, thus without any organic findings or gastrointestinal symptoms, which could have influenced the microbiota composition. Also, the sequencing depth of the biopsy-derived samples (0.15 million unique read-counts) was limited by the large number of human genes sampled together with the microbes. This is another unavoidable limitation of this method. The lower sequencing depth led to a partial measurement of the gut mucosal microbiota, which mainly emphasised the most abundant species. While these are both important and expected limitations, we still chose to use this method with the aim of assessing what can be learnt from it. Biopsies can provide information that faeces cannot: subtle and otherwise undetectable variations in local microbiomes along the intestine that faeces cannot detect. Such variations are of great interest since, as discussed, they can affect the onset and progression of serious pathologies. We thus believe these results, although still not optimal, provide new valuable insights into the mucosal gut microbiota. New methods will need to be devised to better isolate microbial species from human tissues, thus improving the sequencing depth. Also, given the promising results here described, larger cohorts now need to be investigated.

In conclusion, the current study showed that faecal samples provide a good approximation of the gut mucosal microbiome, although only metagenomics of gut mucosal biopsies can detect subtle variation in the local microbial communities’ composition along the large intestine. Functional analysis of the biopsy metagenomics data was in agreement with the current knowledge while providing new fundamental information. Our GEM simulation, in particular, could detail which species are involved in only partially known metabolic mechanisms connected with health or disease. Our work provides the first detailed insight into which microbial species are associated with the gut mucosal microbiota of healthy individuals. Such valuable information will provide the starting point for more targeted future investigations on the gut microbiota.

## Methods

Study population. Five healthy adult volunteers from Stockholm were selected from the participants to a study previously described^25^. Colonoscopy preparation included a clear liquid diet and bowel cleansing with 45mL Phosphoral oral intake twice within 4 hours. At colonoscopy, biopsies were collected from terminal ileum, caecum, transverse colon, and rectum. Faecal samples were collected at home, before bowel cleansing and colonoscopy, and sent by post to the research facility where they were frozen at - 80 °C. All subjects were free from any objective finding at colonoscopy. They had not undergone any previous gastrointestinal surgeries and had no current or previous diseases of the gastrointestinal tract (for exclusion criteria, see Supplementary Materials). The study was approved by the Karolinska Institutet ethical review board (Forskningskommitté Syd, nr 394/01). All participants provided written informed consent.

DNA extraction. Total genomic DNA was isolated from biopsy tissue or from 100-120 mg faecal sample using repeated bead beating, following a protocol previously described^50^. Briefly, samples were placed in Lysing Matrix E tubes (MP Biomedicals), and sterile lysis buffer (4% w/v SDS; 500 mmol/L NaCl; 50 mmol/L EDTA; 50 mmol/L Tris•HCl; pH 8) was added. Biopsy samples were incubated with 50 µl mix of mutanolysin (5U/µL) and lysozyme (100 mg/ml) at 37 °C for 30 min. Both biopsy and faeces samples were lysed twice with bead beating at 5.0 m/s for 60 s in a FastPrep®-24 Instrument (MP Biomedicals). After each bead-beating, samples were heated at 85°C for 15 minutes and centrifuged at full speed for 5 min at 4°C. Supernatants from the two lysate-fractions were pooled and purified. Total genomic DNA was eluted in AE buffer (10 mmol/L Tris•Cl; 0.5 mmol/L EDTA; pH 9.0).

Analysis of shotgun metagenomics. Extracted DNA was processed into a paired-end library and sequenced by Illumina HiSeq 2500 (2×100bp), generating an average of 5.9 million paired-end reads per sample. Raw data was quality checked with FastQC (http://www.bioinformatics.babraham.ac.uk/projects/fastqc), processed with METEOR^51^, and mapped onto the integrated reference catalogue of the human gut microbiome^52^. Host DNA was removed. Aligned reads were counted, normalized, and then downsized by R package MetaOMineR^26^. Downsizing, in particular, was performed in order to take into account the different sequencing depths across samples. Strain/species-level abundances of metagenomic species (MGS) – i.e., co-abundant genes with more than 100 genes originated from microbial species^53, 54^ – were profiled for each sample. Downsizing was performed for the whole biopsy-faeces dataset, then also for biopsy and faeces datasets separately. MGS profiles were estimated as the mean abundance of the 50 genes of a given MGS (centroid of the clustered genes) and used to perform taxonomic investigations. Based on the orthologous genes from the KEGG database^55^, we annotated the quantified bacterial genes, and performed functional analysis of the genes detected in the biopsy dataset.

### Statistical analysis

To assess if and how much each set of data (either sampling location- or patient-specific) differ, in the average, from the others, paired difference tests were performed. In particular, as the available data was not large and did not appear to be normally distributed, statistical assessments were performed with the Wilcoxon signed-rank test^56^, in MatLab.

### Antibiotic resistance genes

All fastq files were mapped against the nucleotide_fasta_protein_homolog_model from the antimicrobial resistance database CARD3.0.0^57^ (https://card.mcmaster.ca/) using Bowtie2 (ver.2.3.4.3). Unmapped reads were filtered out, and resistance genes with mapped reads coverage below 90% were discarded. Each resistance gene was annotated with Drug Class and Resistance Mechanism using CARD3.0.0 metadata. R package Deseq2 was used to normalize the resistome dataset, and to perform statistical evaluation of significantly changing resistance genes.

### Secondary metabolite identification via antiSMASH pipeline

All the gene sequences of the 606 metagenome species we identified were retrieved from the reference gene catalogue^52^, and the antiSMASH standalone program was used to annotate their biosynthetic genes by minimal run options focused on core detection modules (version 5)^58^. The antiSMASH program was loaded onto the Amazon cloud computing platform (AWS) as docker image, and its mining process was executed per metagenomics species with all processes massively parallelized. All detected secondary metabolite clusters per metagenomics species were then associated with the sampling locations of each metagenomics sample.

### Genome Scale metabolic Model (GEM) reconstruction and simulation

We reconstructed the GEMs of the bacteria (Supplementary Table 1) belonging to genera *Parabacteroides, Anaerostipes, Bacteroides, Bifidobacterium, Eubacterium, Ruminococcus*, and *Blautia*, using the KO annotation provided in the gut catalogue. The KO profiles were converted to metabolic network and reaction score profiles regarding the KBase reference model. To make the functional models regarding the provided biomass objective function, the gap filling was done using the raven toolbox. All models were constrained by the general UK diet (https://fdnc.quadram.ac.uk/). Simulations were run anaerobically to calculate the growth rate for each model, and the production profiles of the bacteria. The resulting biomass figures were proportioned by species abundances based on colon locations. We added an initial source of acetate and lactate on the basis of the average production profile of each microbe. For the second simulation, models were constrained both by the predicted biomass of the bacteria and by the diet. The flux balance analysis result (Supplementary Table 2) was used to find metabolite importance for each part of the colon. The sensitivity of the models to oxygen was simulated regarding the oxygen-uptake of the intestine (2 ml O_2_/min*100 grams tissue), where the micro-aerobic condition was considered as 5% O_2_-uptake. All references relative to the GEM method here described are available in Supplementary Materials.

## Data availability

All raw metagenomic data have been deposited in the public EBI/NCBI Database under accession number PRJEB33194.

## Acknowledgements

We acknowledge the support from Science for Life Laboratory, the National Genomics Infrastructure (NGI), and Uppmax for providing assistance in parallel sequencing and computational infrastructure.

## Author contributions

S.V. wrote the manuscript. S.V. and S.L. analysed and interpreted the data. B.J. performed metagenomics data acquisition. L.A. and N.J.T. designed and supervised the origin PopCol study, A.A. supervised the study and worked on all logistics on data entry and outcome. P.K.D. processed all the samples and prepared the DNA libraries for sequencing. G.B. reconstructed the GEM models and simulated the models. J.N., L.E. and S.S. designed the project. D.E. and S.S. provided important critical revisions of the manuscript and helped to interpret the data. D.L. and J.P. performed the secondary metabolite annotation. N.J.T., G.P., D.E., J.N., L.E., and S.S. provided critical revision of the manuscript. All authors contributed to discuss the results and to research directions. All authors approved the manuscript.

## Competing interests

N.J.T. reports personal fees from Allergans PLC (GI Development Programs), personal fees from Viscera Labs (IBS), personal fees from IM Health Sciences (FD), personal fees from Napo Pharmaceutical (IBS), personal fees from Outpost Medicine (IBS), from ProgenityInc San Diego (capsule SIBO), from Allakos (gastric eosinophilic disease), personal fees from Samsung Bioepis (IBD), personal fees from Synergy (IBS), personal fees from Takeda (gastroparesis), personal fees from Theravance (gastroparesis), grants and personal fees from Viscera USA (IBS), grants from Commonwealth Diagnostics (International) Inc (IBS), non-financial support from HVN National Science Challenge NZ (IBS), grants and personal fees from GI therapies (constipation), personal fees from Cadila Pharmaceuticals (CME), personal fees from Planet Innovation (Gas capsule), personal fees from Danone (Probiotic), personal fees from Pfizer (IBS), from Dr. Reddy’s Laboratories (Webinar), personal fees from Arlyx (IBS), personal fees from Sanofi (Probiotic), outside the submitted work; In addition, Dr. Talley has a patent Biomarkers of IBS licensed, a patent Licensing Questionnaires Talley Bowel Disease Questionnaires licensed to Mayo/Talley, a patent Nestec European Patent licensed, a patent Singapore Provisional Patent “Microbiota Modulation Of BDNF Tissue Repair Pathway” issued, and a patent Nepean Dyspepsia Index licensed to Talley copyright and Committees: Australian Medical Council (AMC) [Council Member]; Australian Telehealth Integration Programme; MBS Review Taskforce; NHMRC Principal Committee (Research Committee) Asia Pacific Association of Medical Journal Editors. Boards: GESA Board Member, Sax Institute, Committees of the Presidents of Medical Colleges. Community group: Advisory Board, IFFGD (International Foundation for Functional GI Disorders). Miscellaneous: Avant Foundation (judging of research grants). Editorial: Medical Journal of Australia (Editor in Chief), Up to Date (Section Editor), Precision and Future Medicine, Sungkyunkwan University School of Medicine, South Korea.

## Fundings

S.S. has received the project grants from the Engineering and Physical Sciences Research Council (EPSRC) and co-funded by Biotechnology and Biological Sciences Research Council (project EP/ S001301/1). N.J.T. has received project grant funding from the National Health and Medical Research Council of Australia. L.E. has received project grants from the T. Söderbergs foundation. D.E. received funding from Metagenopolis grant ANR-11-DPBS-0001. P.K.D. is a recipient of fellowships from the CAS President’s International Fellowship Initiative (PIFI; Project No. 2018PB0028) and DICP Outstanding Postdoctoral Foundation (Grant No. 2017YB05). DL and JP were supported by the Bio-Synergy Research Project (2012M3A9C4048758) of the Ministry of Science and ICT through the National Research Foundation.

## References

1. Dethlefsen, L., McFall-Ngai, M. & Relman, D. A. An ecological and evolutionary perspective on humang-microbe mutualism and disease. Nature 449, 811–818 (2007).

2. Zhang, Z. et al. Spatial heterogeneity and co-occurrence patterns of human mucosal-associated intestinal microbiota. ISME J. 8, 881–893 (2014).

3. Mcmahon, T., Zijl, P. C. M. Van & Gilad, A. A. Eating For Two: How Metabolism Establishes Interspecies Cell Host Microbe Interactions in the Gut. 27, 320–331 (2015).

4. Goodrich, J. K. et al. Genetic Determinants of the Gut Microbiome in UK Twins. Cell Host Microbe 19, 731–743 (2016).

5. Donaldson, G. P., Lee, S. M. & Mazmanian, S. K. Gut biogeography of the bacterial microbiota. Nat. Rev. Microbiol. 14, 20–32 (2015).

6. Qin, J. et al. Europe PMC Funders Group Europe PMC Funders Author Manuscripts A human gut microbial gene catalog established by metagenomic sequencing. 464, 59–65 (2013).

7. Vogtmann, E. et al. Colorectal cancer and the human gut microbiome: Reproducibility with whole-genome shotgun sequencing. PLoS One 11, 1–13 (2016).

8. Vila, A. V. et al. Gut microbiota composition and functional changes in inflammatory bowel disease and irritable bowel syndrome. 1–12

9. Kuang, Y. S. et al. Connections between the human gut microbiome and gestational diabetes mellitus. Gigascience 6, 1–12 (2017).

10. Jandhyala, S. M. et al. Role of the normal gut microbiota. World J. Gastroenterol. 21, 8836–8847 (2015).

11. Ottman, N., Geerlings, S. Y., Aalvink, S., Vos, W. M. De & Belzer, C. Best Practice & Research Clinical Gastroenterology Action and function of *Akkermansia muciniphila* in microbiome ecology, health and disease. Best Pract. Res. Clin. Gastroenterol. 31, 637–642 (2017).

12. Lee, H. et al. Modulation of the gut microbiota by metformin improves metabolic profiles in aged obese mice. Gut Microbes 9, 155–165 (2018).

13. Cekanaviciute, E. et al. Gut bacteria from multiple sclerosis patients modulate human T cells and exacerbate symptoms in mouse models. Proc. Natl. Acad. Sci. 114, 201711235 (2017).

14. Bedarf, J. R. et al. Functional implications of microbial and viral gut metagenome changes in early stage L-DOPA-naïve Parkinson’s disease patients. Genome Med. 9, 1–13 (2017).

15. Ghotaslou, R., Yekani, M. & Memar, M. Y. The role of e ffl ux pumps in bacteroides fragilis resistance to antibiotics. Microbiol. Res. 210, 1–5 (2018).

16. Cirkeline, K. et al. Anaerobe Antimicrobial resistance in the bacteroides fragilis group in faecal samples from patients receiving broad-spectrum antibiotics. Anaerobe 47, 79–85 (2017).

17. Deng, H. et al. Bacteroides fragilis Prevents Clostridium difficile Infection in a Mouse Model by Restoring Gut Barrier and Microbiome Regulation. 9, 1–12 (2018).

18. Mazmanian, S. K., Liu, C. H., Tzianabos, A. O. & Kasper, D. L. of Symbiotic Bacteria Directs Maturation of the Host Immune System. 122, 107–118 (2005).

19. Sommer, F. & Bäckhed, F. The gut microbiota-masters of host development and physiology. Nat. Rev. Microbiol. 11, 227–238 (2013).

20. Beaumont, M. et al. Quantity and source of dietary protein influence metabolite production by gut microbiota and rectal mucosa gene expression: A randomized, parallel, double-blind trial in overweight humans. Am. J. Clin. Nutr. 106, 1005–1019 (2017).

21. Howell, K. J. et al. DNA Methylation and Transcription Patterns in Intestinal Epithelial Cells From Pediatric Patients With Inflammatory Bowel Diseases Differentiate Disease Subtypes and Associate With Outcome. Gastroenterology 154, 585–598 (2018).

22. Li, G. et al. Diversity of duodenal and rectal microbiota in biopsy tissues and luminal contents in healthy volunteers. J. Microbiol. Biotechnol. 25, 1136–1145 (2015).

23. Suez, J. et al. Post-Antibiotic Gut Mucosal Microbiome Reconstitution Is Impaired by Probiotics and Improved by Autologous FMT. Cell 174, 1406–1423.e16 (2018).

24. Hillmann, B. et al. Evaluating the Information Content of Shallow Shotgun Metagenomics. 3, 1–12 (2018).

25. Kjellström, L. et al. A randomly selected population sample undergoing colonoscopy: Prevalence of the irritable bowel syndrome and the impact of selection factors. Eur. J. Gastroenterol. Hepatol. 26, 268–275 (2014).

26. Le Chatelier, E. et al. Richness of human gut microbiome correlates with metabolic markers. Nature 500, 541–546 (2013).

27. Leite, A. Z. et al. Detection of increased plasma interleukin-6 levels and prevalence of Prevotella copri and *Bacteroides* vulgatus in the feces of type 2 diabetes patients. Front. Immunol. 8, (2017).

28. Chiodini, R. J. et al. Transitional and temporal changes in the mucosal and submucosal intestinal microbiota in advanced crohn’s disease of the terminal ileum. J. Med. Microbiol. 67, 549–559 (2018).

29. Chiodini, R. J., Dowd, S. E., Galandiuk, S., Davis, B. & Glassing, A. The predominant site of bacterial translocation across the intestinal mucosal barrier occurs at the advancing disease margin in Crohn’s disease. Microbiol. (United Kingdom) 162, 1608–1619 (2016).

30. Øyri, S. F., Muzes, G. & Sipos, F. Dysbiotic gut microbiome: A key element of Crohn’s disease. Comp. Immunol. Microbiol. Infect. Dis. 43, 36–49 (2015).

31. van Schaik, W. The human gut resistome. Philos. Trans. R. Soc. B Biol. Sci. 370, (2015).

32. Kanehisa, M., Sato, Y., Kawashima, M., Furumichi, M. & Tanabe, M. KEGG as a reference resource for gene and protein annotation. Nucleic Acids Res. 44, D457–D462 (2016).

33. Lyu, Y., Wu, L., Wang, F., Shen, X. & Lin, D. Carotenoid supplementation and retinoic acid in immunoglobulin A regulation of the gut microbiota dysbiosis. Exp. Biol. Med. 243, 613–620 (2018).

34. Medema, M. H. et al. AntiSMASH: Rapid identification, annotation and analysis of secondary metabolite biosynthesis gene clusters in bacterial and fungal genome sequences. Nucleic Acids Res. 39, 339–346 (2011).

35. Schöner, T. A. et al. Aryl Polyenes, a Highly Abundant Class of Bacterial Natural Products, Are Functionally Related to Antioxidative Carotenoids. ChemBioChem 17, 247–253 (2016).

36. Cummings, J. H., Pomare, E. W., Branch, H. W. J., Naylor, C. P. E. & MacFarlane, G. T. Short chain fatty acids in human large intestine, portal, hepatic and venous blood. Gut 28, 1221–1227 (1987).

37. Gao, J. et al. Impact of the Gut Microbiota on Intestinal Immunity Mediated by Tryptophan Metabolism. 8, 1–22 (2018).

38. Wexler, H. M. *Bacteroides*: The good, the bad, and the nitty-gritty. Clin. Microbiol. Rev. 20, 593–621 (2007).

39. Tzianabos, A. O. et al. Structural Features of Polysaccharides That Abscesses Induce Intra-Abdominal. 262, 416–419 (2014).

40. An, D., Na, C., Bielawski, J., Hannun, Y. A. & Kasper, D. L. Membrane sphingolipids as essential molecular signals for *Bacteroides* survival in the intestine. Proc. Natl. Acad. Sci. 108, 4666–4671 (2011).

41. Maskell, J. P. The resolution of bacteroides lipopolysaccharides by polyacrylamide gel electrophoresis. J. Med. Microbiol. 34, 253–257 (1991).

42. Jacobson, A. N., Choudhury, B. P. & Fischbach, M. A. The Biosynthesis of Lipooligosaccharide from *Bacteroides* thetaiotaomicron. MBio 9, 1–14 (2018).

43. Goodman, A. L. et al. Identifying Genetic Determinants Needed to Establish a Human Gut Symbiont in Its Habitat. Cell Host Microbe 6, 279–289 (2009).

44. Pumbwe, L., Skilbeck, C. A. & Wexler, H. M. The Bacteroides fragilis cell envelope: Quarterback, linebacker, coach-or all three? Anaerobe 12, 211–220 (2006).

45. Halstead, J. M. et al. The effects of micronutrient deficiencies on bacterial species from the human gut microbiota. Science (80-.). 347, 1367–1671 (2015).

46. Kau, A. L., Ahern, P. P., Griffin, N. W., Goodman, A. L. & Gordon, J. I. Human nutrition, the gut microbiome and the immune system. Nature 474, 327–336 (2011).

47. Roager, H. M. & Licht, T. R. Microbial tryptophan catabolites in health and disease. Nat. Commun. 9, 1–10 (2018).

48. Agus, A., Planchais, J. & Sokol, H. Review Gut Microbiota Regulation of Tryptophan Metabolism in Health and Disease. Cell Host Microbe 23, 716–724 (2018).

49. Nikolaus, S. et al. Increased Tryptophan Metabolism Is Associated With Activity of Inflammatory Bowel Diseases. Gastroenterology 153, 1504–1516.e2 (2017).

50. Salonen, A. et al. Comparative analysis of fecal DNA extraction methods with phylogenetic microarray: Effective recovery of bacterial and archaeal DNA using mechanical cell lysis. J. Microbiol. Methods 81, 127–134 (2010).

51. Doré, J. et al. Dietary intervention impact on gut microbial gene richness. Nature 500, 585–588 (2013).

52. Wen, C. et al. Quantitative metagenomics reveals unique gut microbiome biomarkers in ankylosing spondylitis. Genome Biol. 18, 1–13 (2017).

53. Nielsen, H. B. et al. Identification and assembly of genomes and genetic elements in complex metagenomic samples without using reference genomes. Nat. Biotechnol. 32, 822–828 (2014).

54. Le Chatelier, E. et al. Gut microbiome influences efficacy of PD-1–based immunotherapy against epithelial tumors. Science (80-.). 359, 91–97 (2017).

55. Kanehisa, M., Sato, Y., Furumichi, M., Morishima, K. & Tanabe, M. New approach for understanding genome variations in KEGG. Nucleic Acids Res. 47, D590–D595 (2019).

56. Forrester, J. & Ury, H. The Signed-Rank (Wilcoxon) test in the rapid analysis of biological data. Lancet 1, 239–41 (1969).

57. McArthur, A. G. et al. The comprehensive antibiotic resistance database. Antimicrob. Agents Chemother. 57, 3348–3357 (2013).

58. Blin, K. et al. AntiSMASH 4.0 - improvements in chemistry prediction and gene cluster boundary identification. Nucleic Acids Res. 45, W36–W41 (2017).

