## Supplementary material for "Compositional and functional differences of the mucosal microbiota along the intestine of healthy individuals"

### Supplementary Materials

#### Study population exclusion criteria:

- Organic findings at the endoscopy
- Previous GI surgery
- Previous or current GI disease or consultations due to GI symptoms
- Known lactose intolerance
- Current use of medicines or current diagnoses
- Antibiotic use previous 6 months
- Smoking
- Snuff use
- Self-reports of abdominal pain, diarrhoea or constipation on the ASQ or Rome II questionnaire
- Overweight
- High blood pressure
- Aberrant blood test (including C-reactive protein and glucose)

#### Study references for Genome Scale metabolic Modelling:

Arkin AP, Cottingham RW, Henry CS, et al. KBase: The United States Department of Energy Systems Biology Knowledgebase. Nat Biotechnol 2018;36:566–569.

Agren R, Liu L, Shoaie S, et al. The RAVEN Toolbox and Its Use for Generating a Genome-scale Metabolic Model for Penicillium chrysogenum. PLoS Comput Biol 2013;9.

Bidkhori G, Benfeitas R, Klevstig M, et al. Metabolic network-based stratification of hepatocellular carcinoma reveals three distinct tumor subtypes. Proc Natl Acad Sci 2018;115:E11874–E11883.

Lutz J, Henrich H, Bauereisen E. Oxygen supply and uptake in the liver and the intestine. Pflügers Arch Eur J Physiol 1975;360:7–15.

Sigalevich P, Cohen Y. Oxygen-dependent growth of the sulfate-reducing bacterium Desulfovibrio oxyclinae in coculture with Marinobacter sp. strain MB in an aerated sulfate-depleted chemostat. Appl Environ Microbiol 2000;66:5019–5023.

**Supplementary Table 1** – Microbial species used for genome scale metabolic modelling. The table file is a separate supplementary file.

**Supplementary Table 2** – Genome scale metabolic model information and flux balance analysis results. The table file is a separate supplementary file.

**Supplementary Table 3** – Study cohort.

| Patient ID | Age | Sex |
|---|---|---|
| **P1** | 34 | F |
| **P2** | 64 | M |
| **P3** | 35 | M |
| **P4** | 31 | M |
| **P5** | 42 | M |

**Supplementary Table 4** – List of all the MGSs that were detected in our samples. The table file is a separate supplementary file.

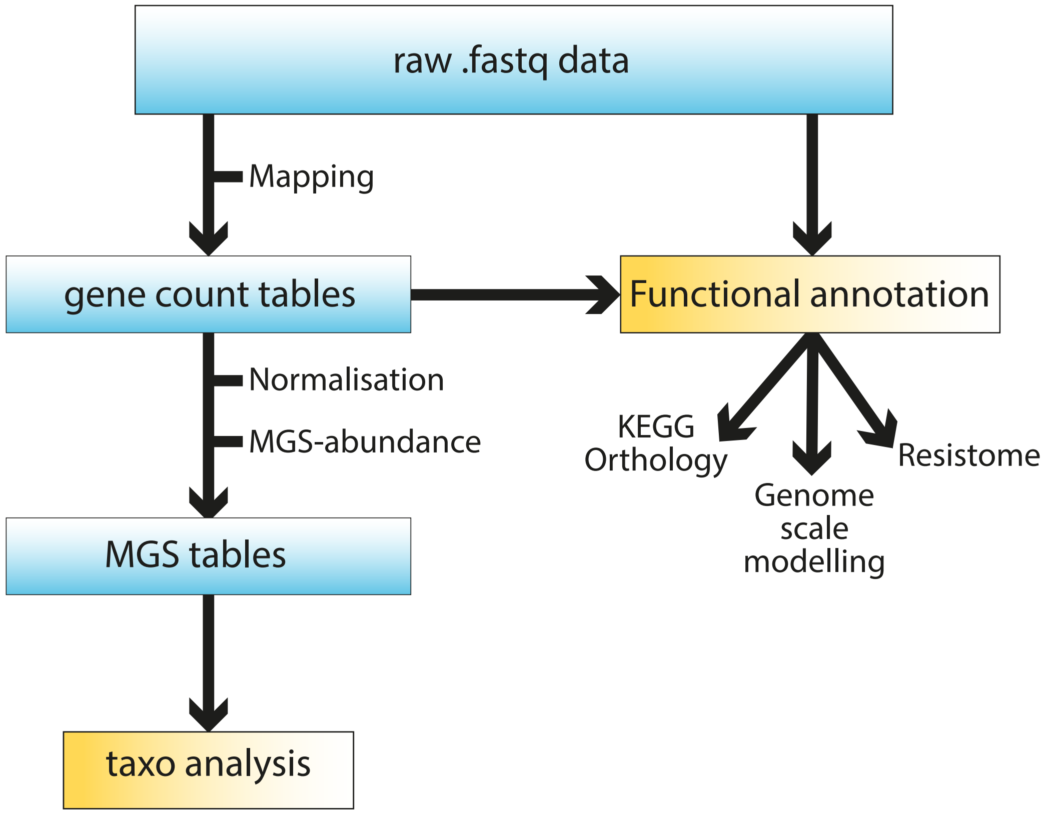


Supplementary Figure 1 - Data analysis workflow for taxonomic and functional investigation of the biopsy/faeces metagenomics data.

A

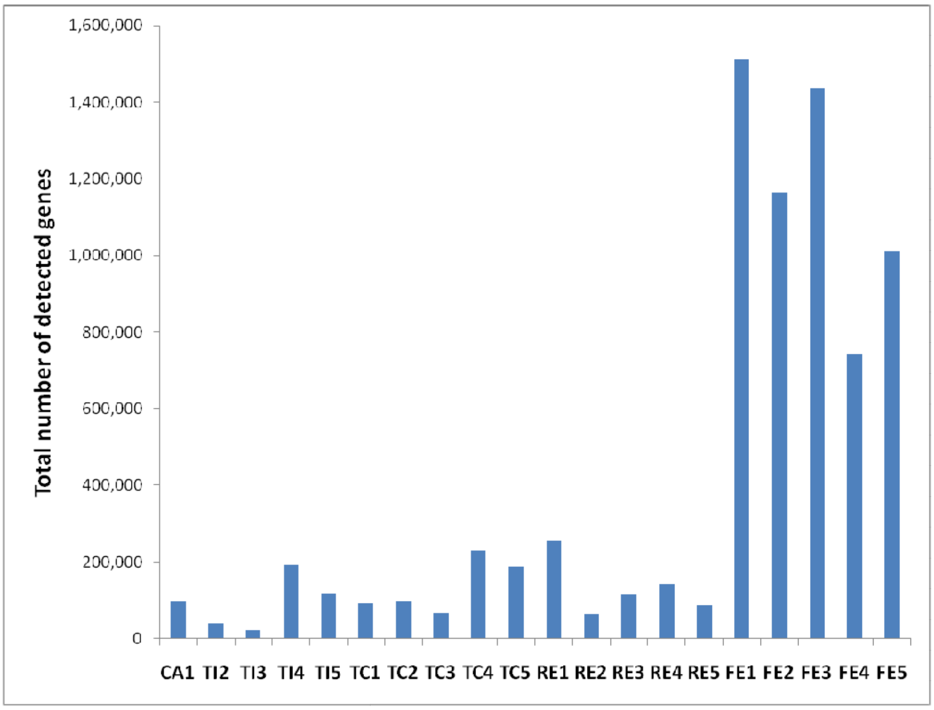


B

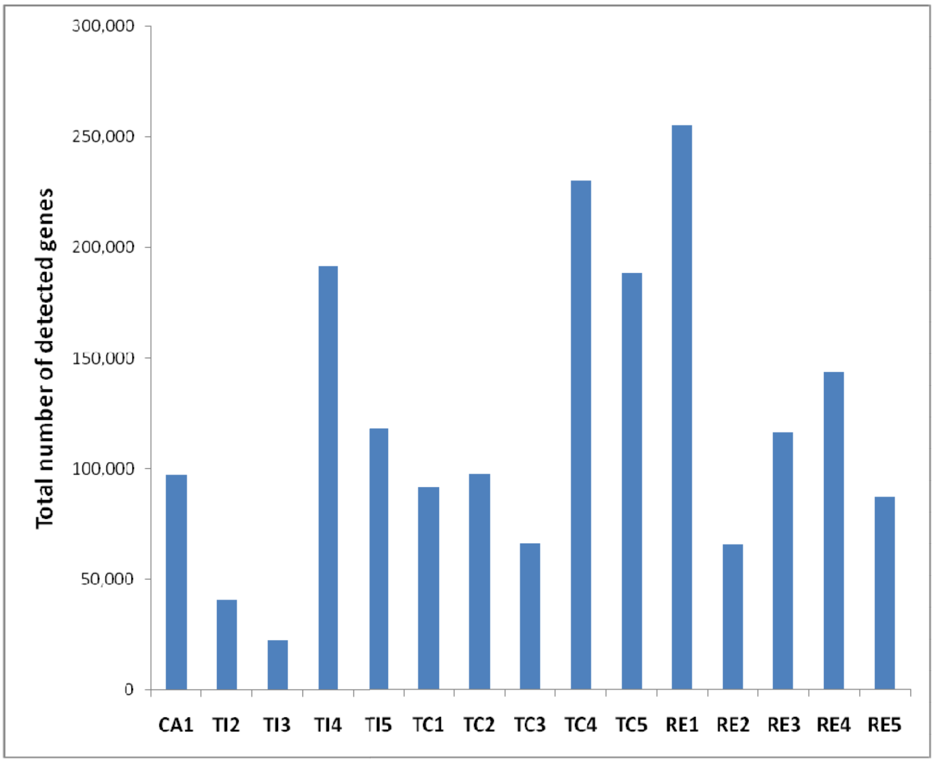


Supplementary Figure 2 - Total number of detected genes in each sample for the complete biopsy plus faeces dataset (A) and for the biopsies only (B).

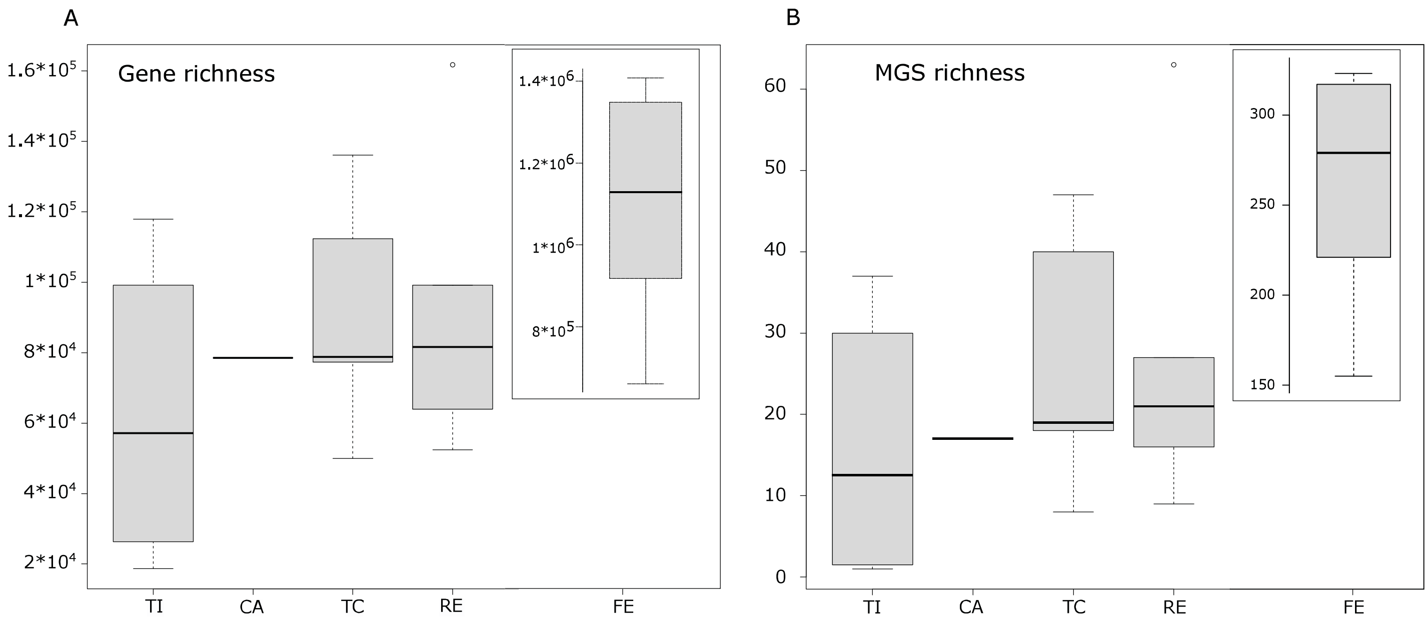


Supplementary Figure 3 - Gene- (A) and MGS- (B) richness boxplots.

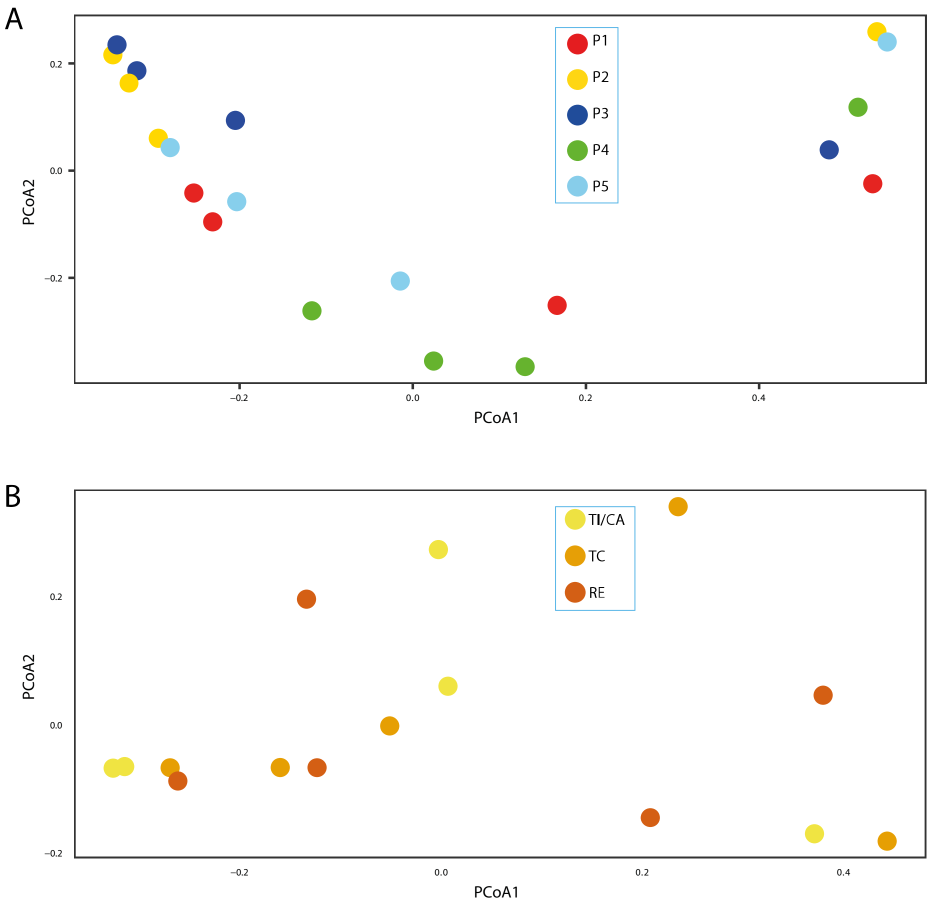


Supplementary Figure 4 - PCoA plots of the complete faeces/biopsies dataset colour-coded for subject (A), and of the biopsies dataset only colour-coded for biopsy sampling location.

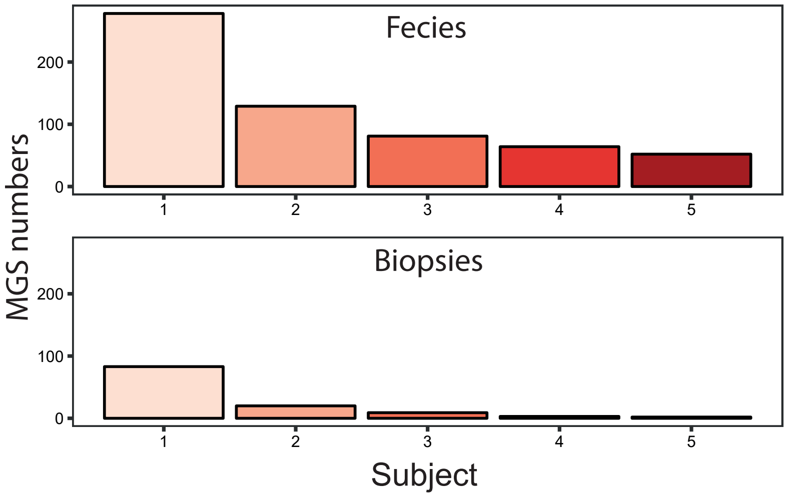


Supplementary Figure 5 - Number of metagenomics species (MGSs) shared between subjects in the faeces dataset (top panel) or in the biopsies one (bottom panel).

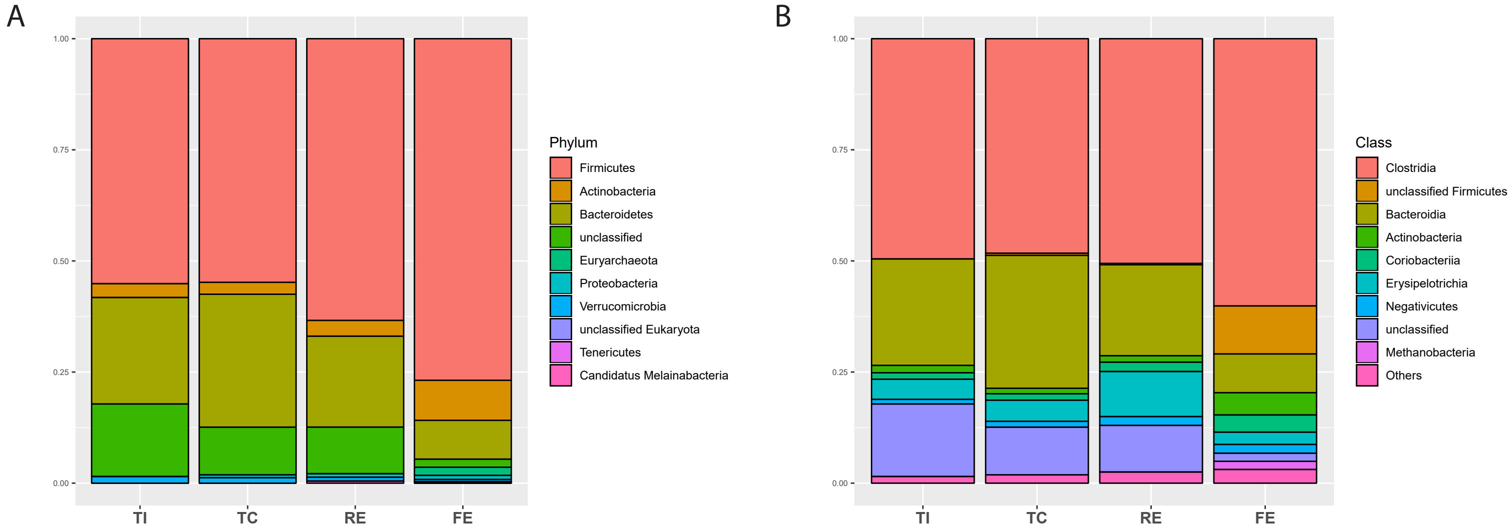


Supplementary Figure 6 - Top 10 most highly abundant phyla (A) and classes (B) in the large intestine; phyla/classes are sorted, in the legends, from the most to the least abundant in the feces samples.

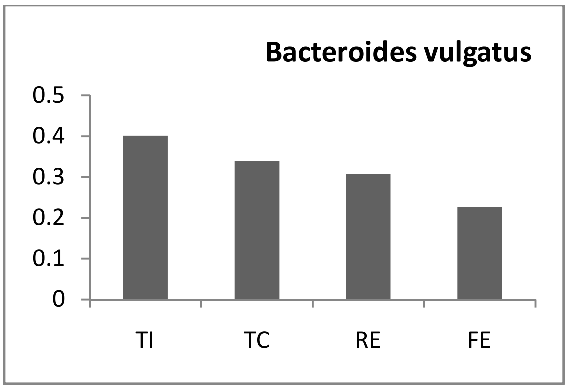
Bacteroides vulgatus
0.5
0.4
0.3
0.2
0.1
0
TI
TC
RE
FE

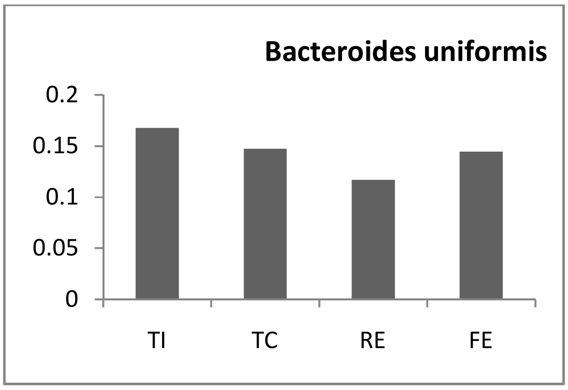
Bacteroides uniformis
0.2
0.15
0.1
0.05
0
TI
TC
RE
FE

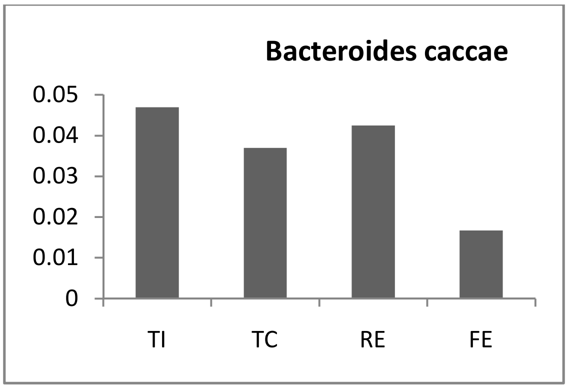
Bacteroides caccae
0.05
0.04
0.03
0.02
0.01
0
TI
TC
RE
FE

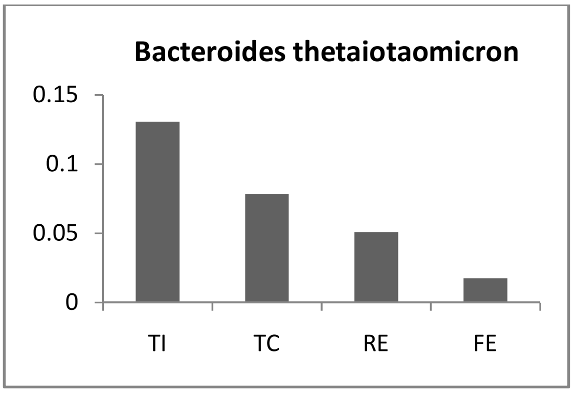
Bacteroides thetaiotaomicron
0.15
0.1
0.05
0
TI
TC
RE
FE

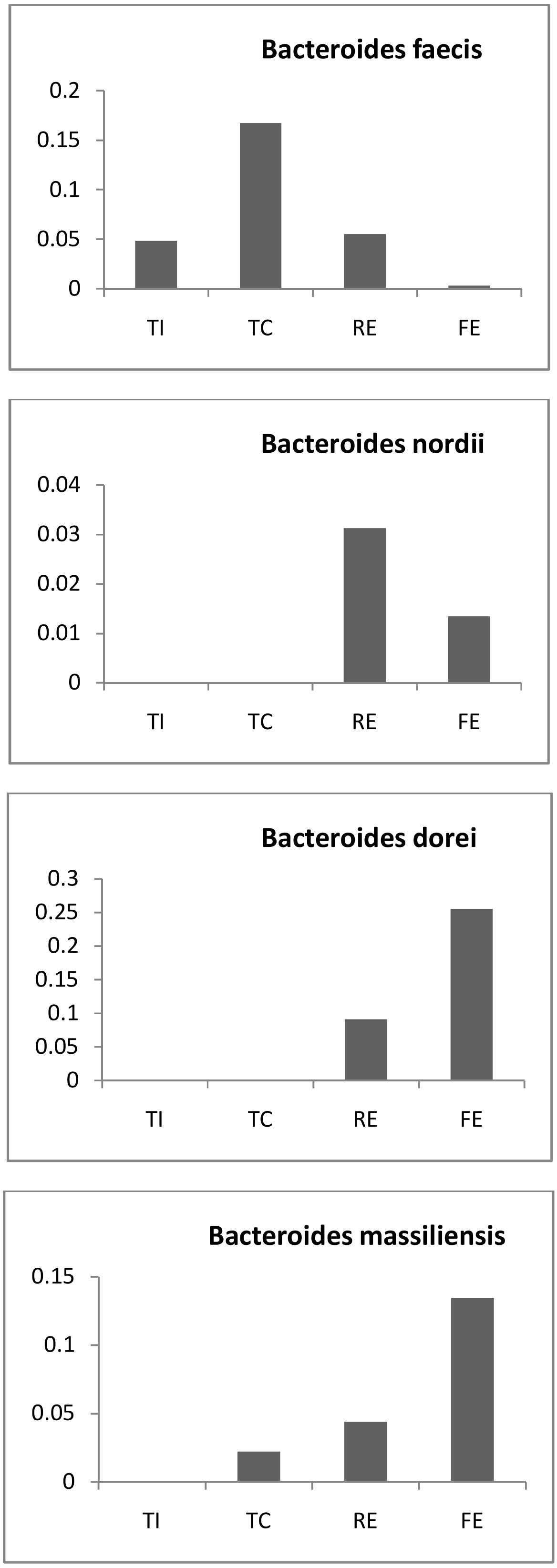


Supplementary figure 7 - Bar plots of the main Bacteroidia detected in the biopsies samples.

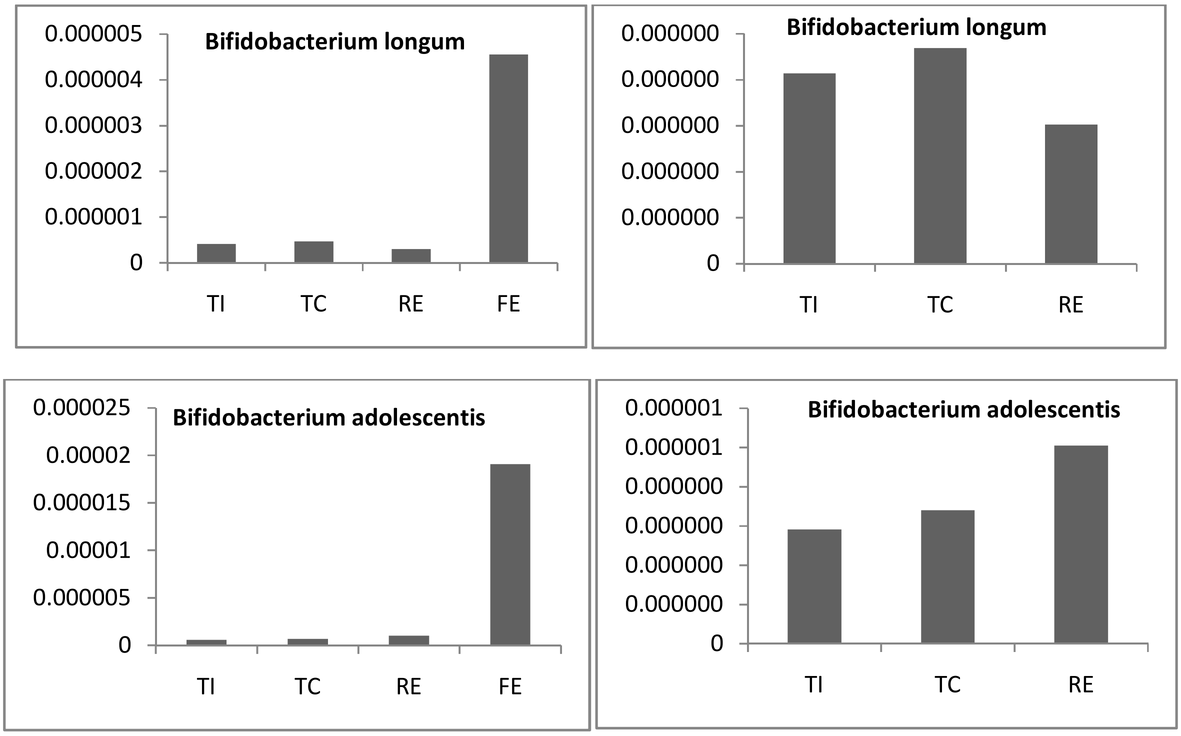


Supplementary figure 8 - Bar plots of the main Bifidobacterium detected in the biopsies and faecal samples.

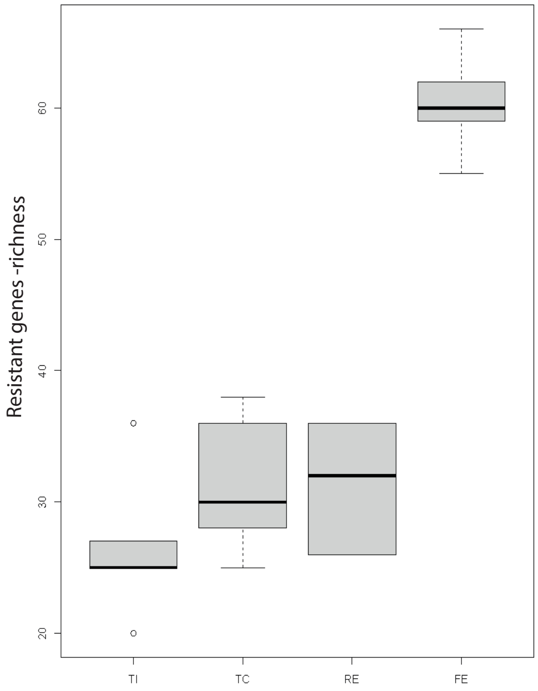


Supplementary figure 9 - Antimicrobial resistant genes - richness in all sampling locations.

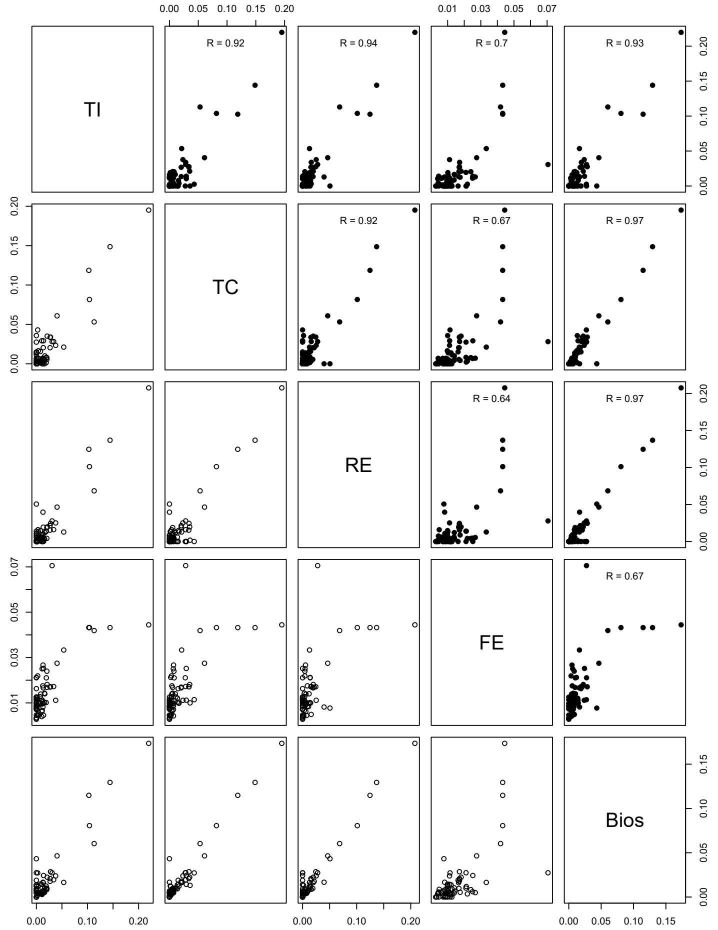


Supplementary figure 10 - Linear correlation between all resistomes.

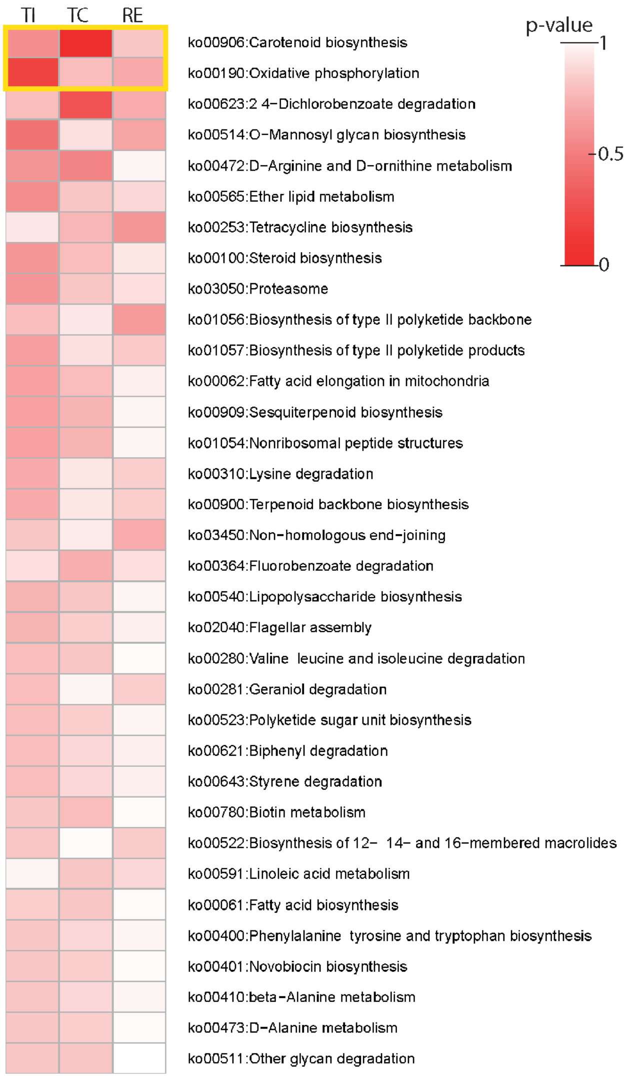


Supplementary Figure 11 - KO orthology by location, results sorted by p-value (Wilcoxon signed rank test). The only two significant pathways are highlighted in yellow.

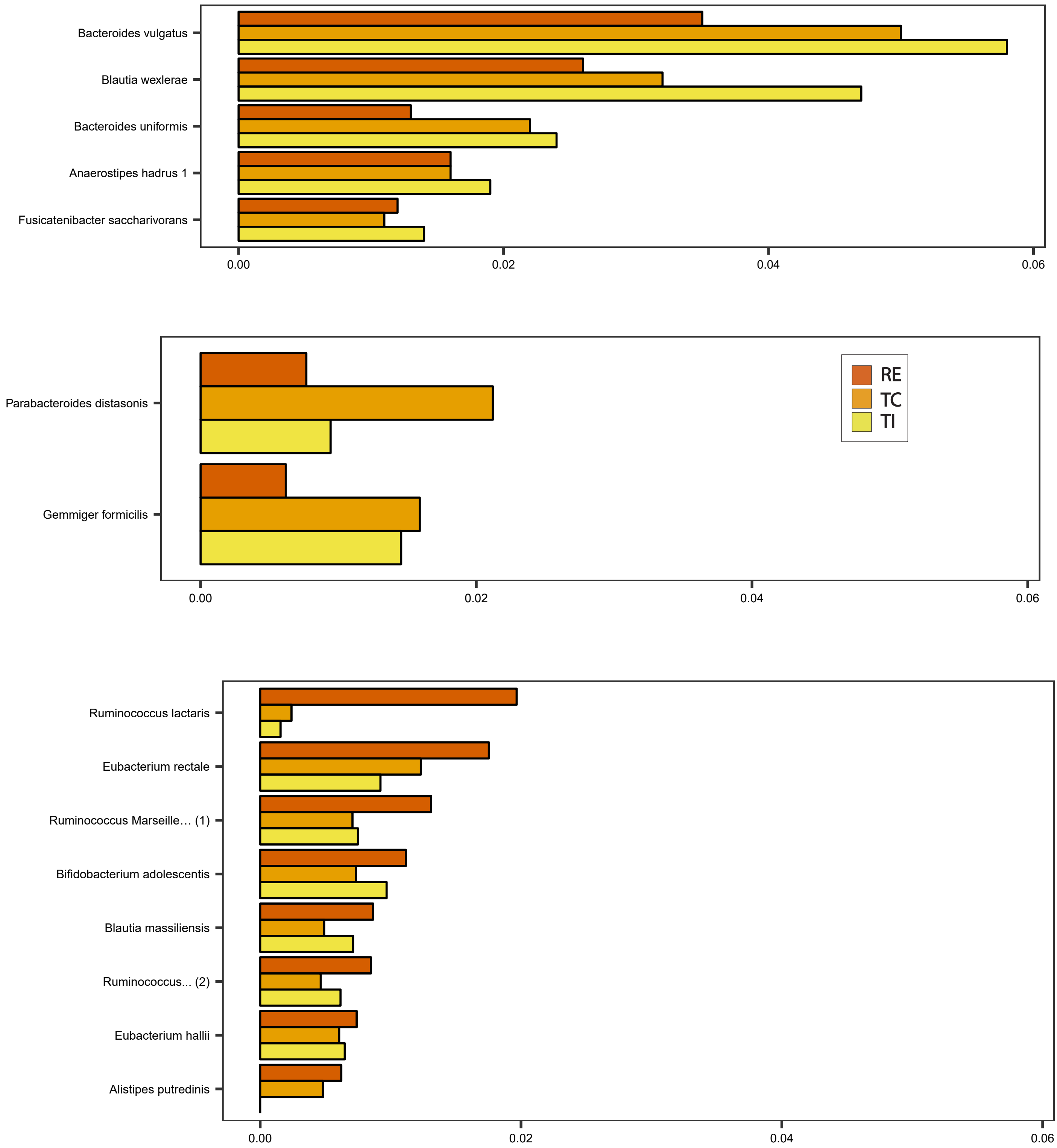


Supplementary figure 12 - Species that were most highly enriched at one biopsy location only, and in at least two patients

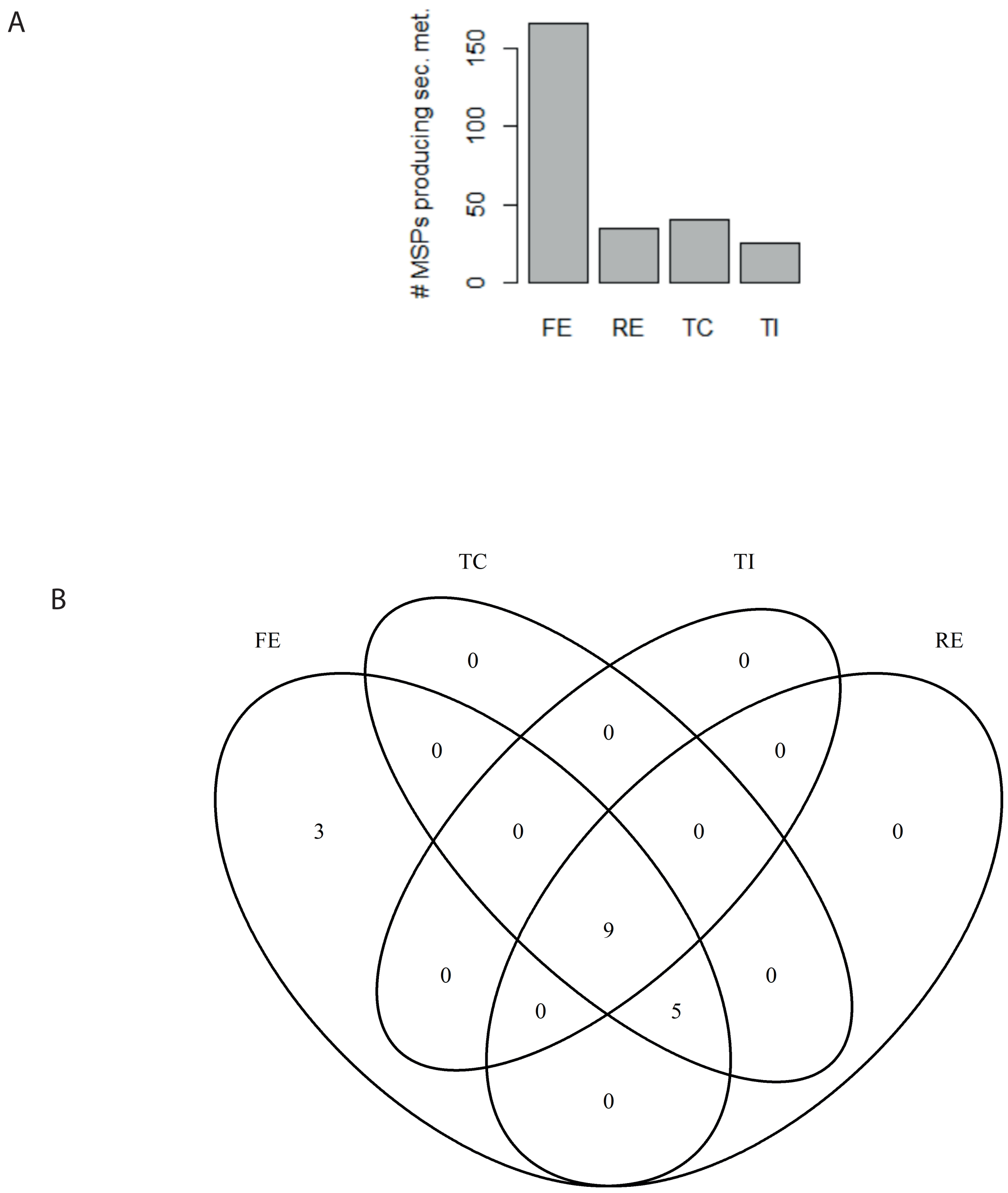


Supplementary Figure 13 - (A) Number of MSPs found to produce secondary metabolites in faeces and biopsy-derived samples. (B) Number of secondary metabolites being shared between different samples.
